## Supplemental Figures for "Single cell transcriptomic profiling of tauopathy in a novel 3D neuron-astrocyte coculture model"

**Author Affiliations**

Benjamin Wolozin

6. Center for Systems Neuroscience, Boston University

Maria Medalla and Benjamin Wolozin

7. Informatics Group, J. Craig Venter Institute, La Jolla, CA, 92037

Dipan Shaw and Christine S. Cheng

8. Department of Psychiatry, University of California San Diego, La Jolla, CA, 92093

Christine S. Cheng

\# These authors contributed equally.

**Supplemental Fig 1**

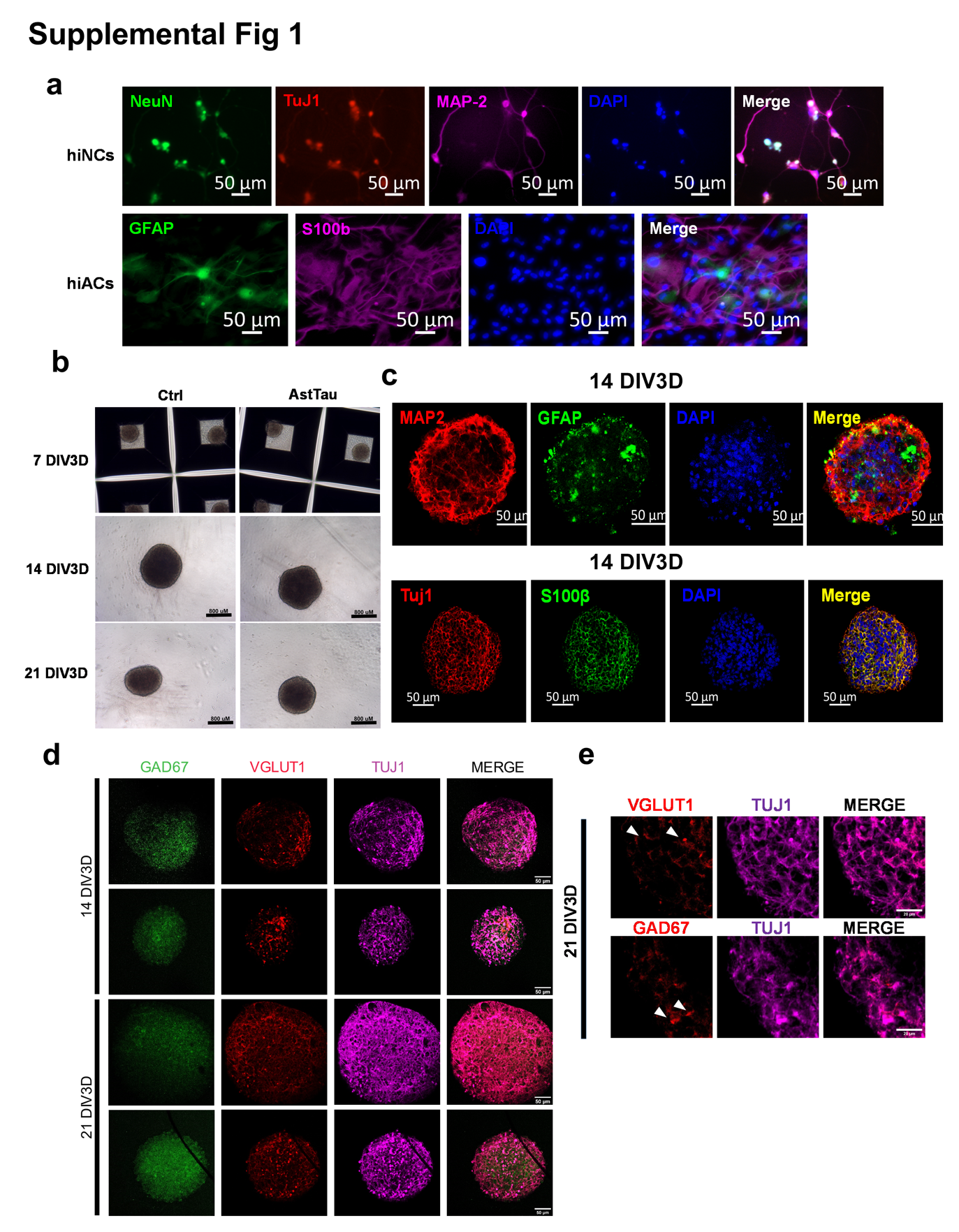

**Supplemental Figure 1. Asteroid cultures are composed of differentiated hiNCs and hiACs that are combined to form self-aggregated spheroidal cultures.**

a. Representative images showing 2D iPSC derived neurons (hiNCs) labeled by NeuN (green), Tuj1 (βIII tubulin, red), MAP2 (violet), and DAPI (blue) and iPSC derived astrocytes (hiACs) labeled by GFAP (green), S100β (violet), and DAPI (blue) at 0 DIV3D, immediately before asteroid generation. Scale bars = 50 µm.

b. Representative images of asteroid cultures at 7, 14, and 21 DIV3D. At 7 DIV3D asteroids are shown cultured in Aggrewell^TM^ microwells. At 14 and 21 DIV3D asteroids are shown cultured in 96 well protein lo-bind wells (see Methods). = 800 µm.

c. Representative images of asteroid cultures at 14 DIV3D. Analysis of cell type markers showed robust differentiation of neurons that were positive for MAP2 and Tuj1 (red) and astrocytes that were positive for S100β and GFAP (green). Scale bars = 50 µm.

d. Representative images of asteroid cultures at 14 and 21 DIV3D showed the presence of inhibitory neuron marker GAD67 (green) and the excitatory neuron marker VGLUT1 (red), which are co-localized in part to the Tuj1 (magenta) neurons, respectively. Scale bars = 50 µm.

e. Representative images of high magnification to show the presence of GAD67 and VGLUT1 in Tuj1+ neurons of 21 DIV3D asteroids. Scale bars = 20 µm.

**Supplemental Fig 2**

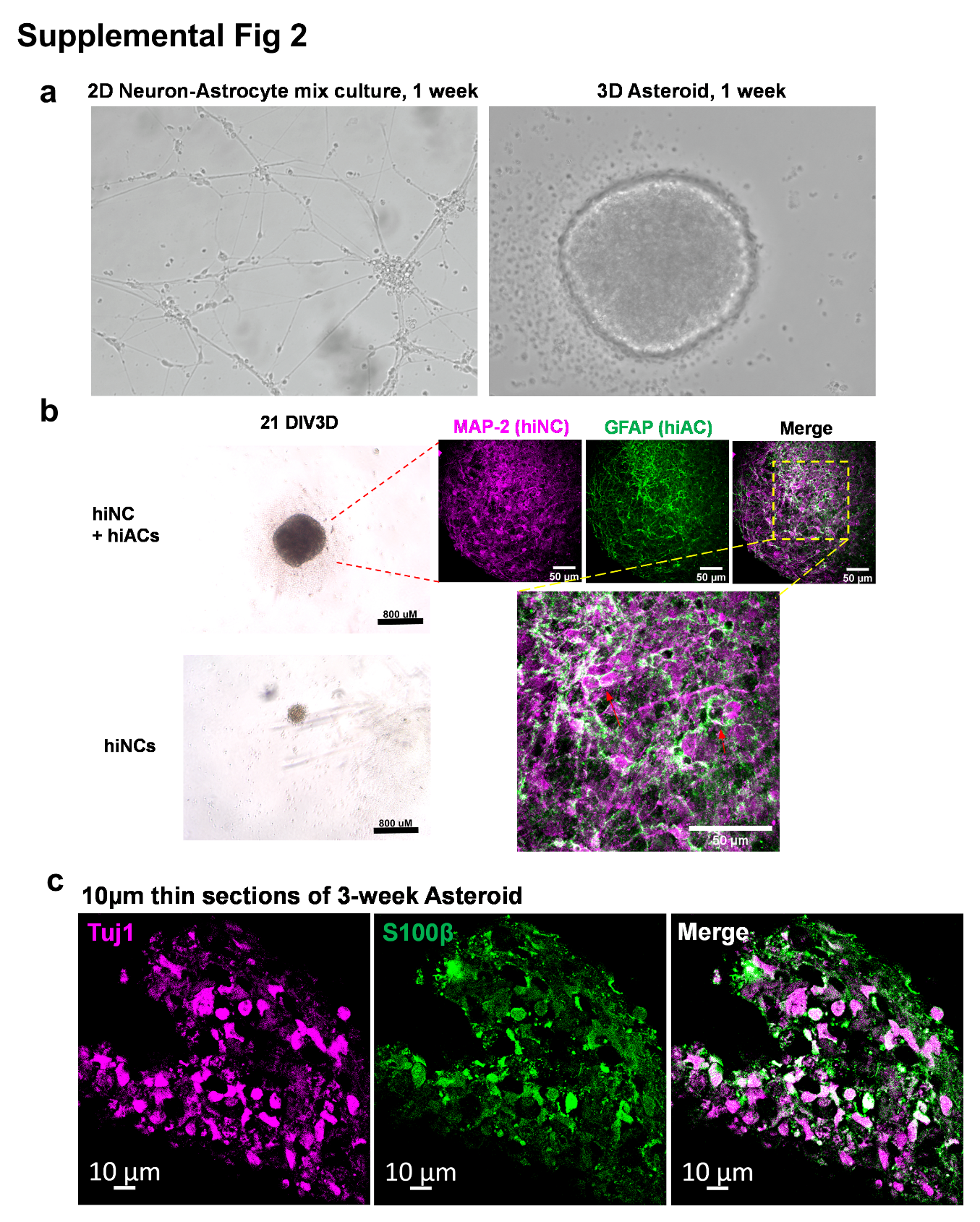

**Supplemental Figure 2. 3D asteroid cultures are characterized by close interactions of hiNCs and hiACs.**

a. Comparison of 2D and 3D neuron-astrocyte co-culture. The 3D asteroid co-culture presented more comprehensive and dimensional interactions of neurons and astrocytes at 1 week.

b. Representative images demonstrating the importance of hiAC incorporation for asteroid generation and survival. Top image of an asteroid composed of hiNCs and hiACs while bottom image of a spheroidal culture composed of hiNCs only, at 21 DIV3D. Immuno-labeling of hiNCs (by MAP-2 antibody, magenta) and hiACs (by GFAP antibody, green) showed the intimate association of neuronal and astrocytic processes in 3D asteroid (red arrows). Scale bars = 50 µm.

c. Representative images of asteroid in high resolution by 10µm sections. This high-resolution imaging shows the intimate proximity of the neurons (Tuj1) and astrocytes (S100β), which complements the extensive arborization and interactions shown in b. Scale bars = 10 µm.

**Supplemental Fig 3**

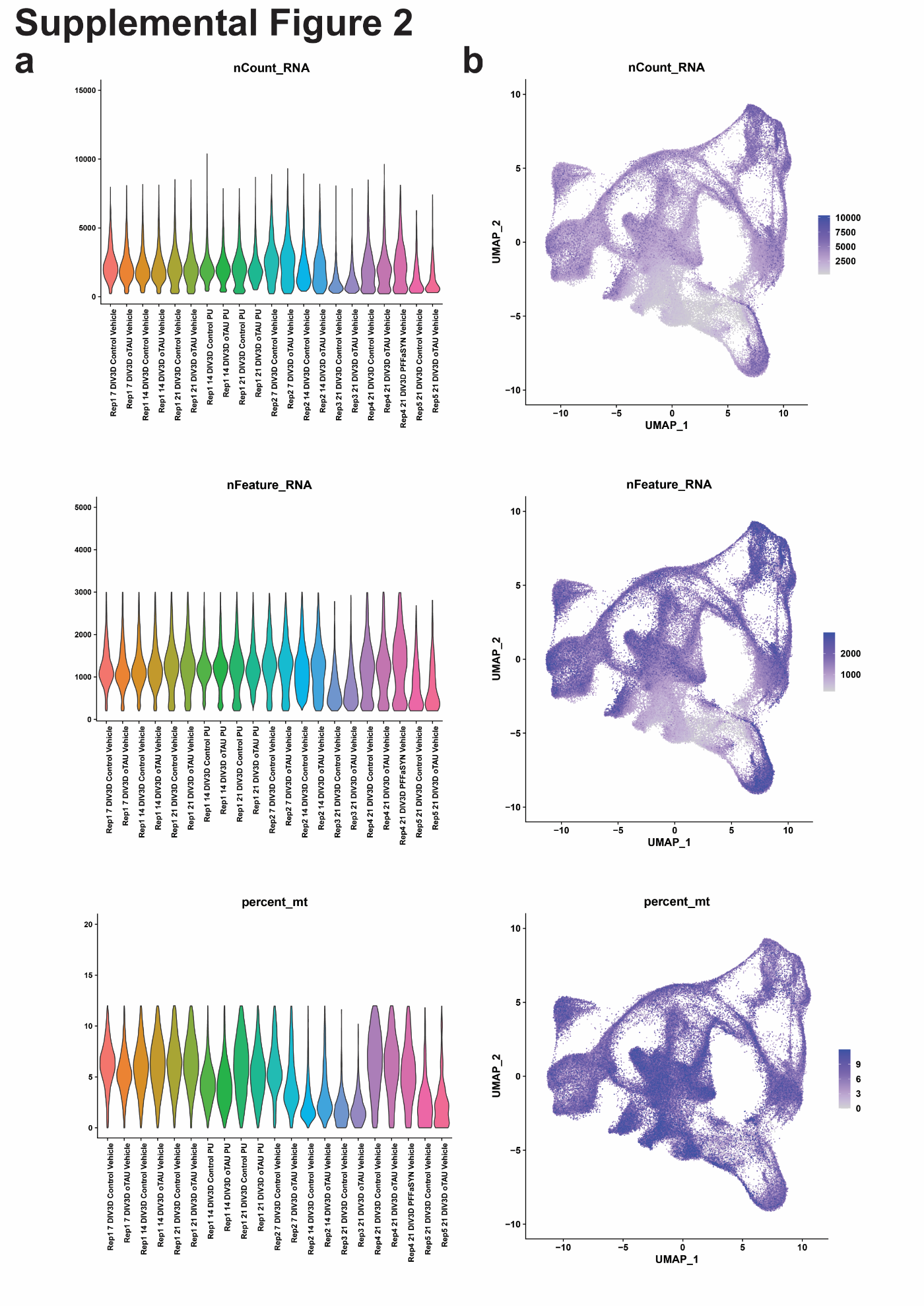

**Supplemental Figure 3. Single cell RNA sequencing captured 130,605 cell transcriptomes across 5 replicate asteroid culture batches and 10 experimental conditions.**

a. Library quality metrics including UMI counts per sample (nCount_RNA), genes per sample (nFeature_RNA), and percent mitochondrial transcripts per sample (percent_mT).
b. UMAP feature plot representation of library quality metrics including UMI counts per cell (nCount_RNA), genes per cell (nFeature_RNA), and percent mitochondrial transcripts per cell (percent_mT).

**Supplemental Fig 4**

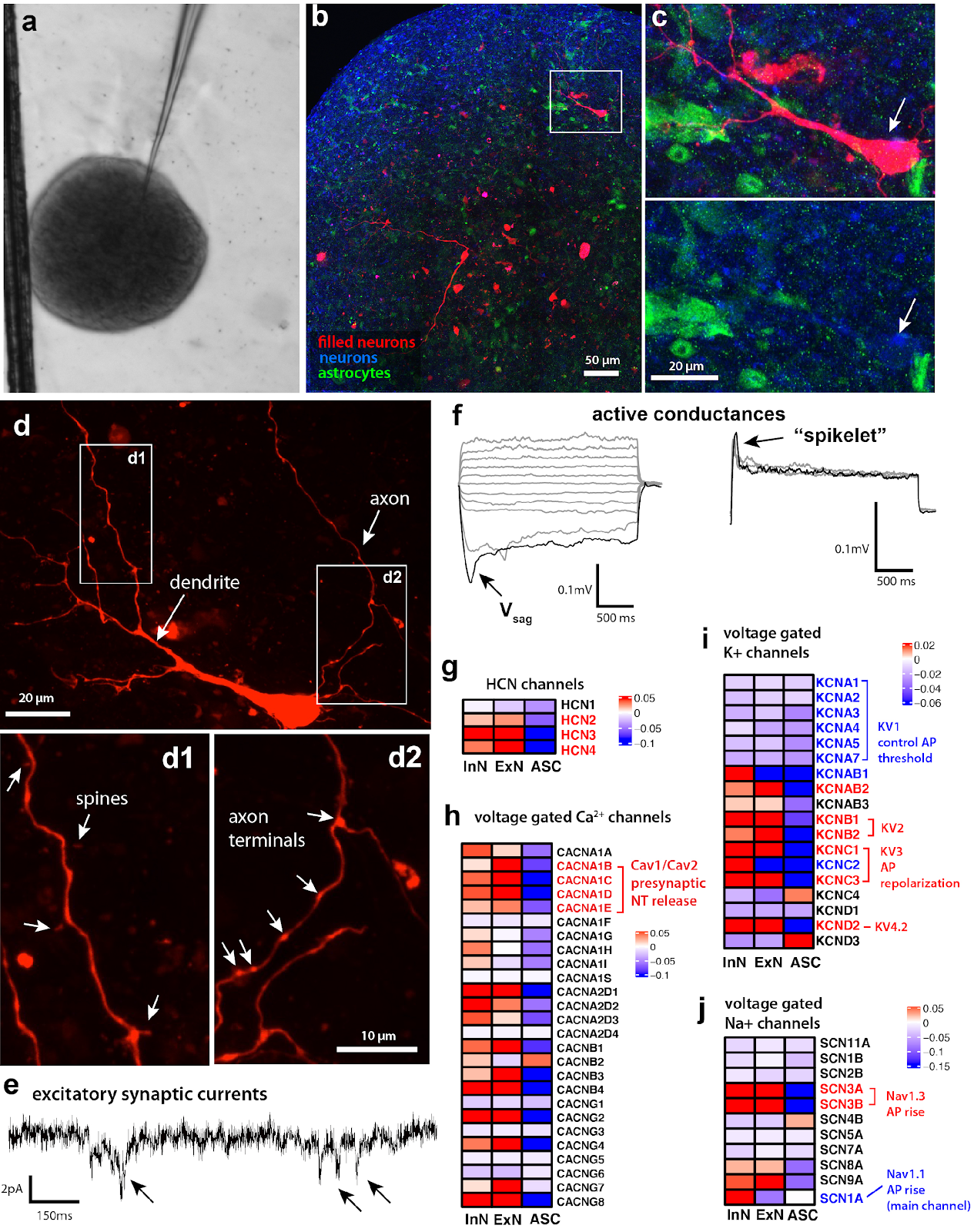

**Supplemental Figure 4. Electrophysiological profiling of asteroid neurons via whole-cell patch-clamp recording and intracellular filling.**

a. Brightfield image of asteroid and glass patch electrode for whole-cell patch-clamp single cell recording and intracellular filling.

b. Fluorescence confocal image montage of Asteroid containing two recorded and filled neurons (red), and immunolabeled for neuronal (TUJ1, blue) and astrocyte (S100B, green) markers to validate cellular phenotype of recorded cells.

c. High magnification 3-channel and green/blue merged images of inset in b, showing a recorded and filled neuron (red, arrow), confirmed to be co-labeled with neuronal marker, TUJ1 (blue), and negative for astrocytic marker S100B (green).

d. High magnification image of filled neuron showing dendrites with spines (d1), and axon with visible axon terminals/boutons (d2).

e. Representative trace of excitatory postsynaptic currents (arrows) recorded under voltage clamp (V_hold_ -80mV).

f. Left, representative traces of voltage responses in response to a series of hyperpolarizing and depolarizing 2sec current steps. Note the presence of a depolarizing voltage sag (V_sag_), in response to hyperpolarizing currents (-100pA), but no evidence of action potentials in response to depolarizing current steps (+80pa to +180pA); Right, high amplitude 2-sec current injections (+220pA, +280pA, +330pA) elicited a depolarizing hump or spikelet voltage response.

g-j. Normalized gene expression profiles of channels implicated in neuronal biophyisical responses single-cell RNA sequencing: g) HCN channel subunits responsible for the Ih current that elicits a depolarizing sag potential (as seen in panel f); h) Voltage-gated Ca2+ channel expression, showing high expression of genes for Cav1 and Cav2 subunits in excitatory neurons (ExN) which are critical for neurotransmitter release. i) Gene expression level of some key voltage gated potassium (K) channels (note that some genes are not shown) reveals a low expression in ExN of genes for KV1 subunits- the low threshold K+ channels that control AP threshold. Apart from the KV2 and KV3 channel genes, which are well expressed, other K channel subunits (not shown) are also not enriched in the asteroid ExN. j)  Voltage gated sodium (Na) channel gene profile shows low expression of the Nav1 subunit gene, which is the main Na channel carrying majority of the sodium current for AP rising phase. Note the relatively high expression of Nav3 channel subunits, which partially contribute to the AP rise sodium current, albeit to a lesser degree than the Nav1 channels. Abbreviations: ExN, excitatory neurons; InN, inhibitory neurons; ASC, astrocytes

**Supplemental Fig 5**

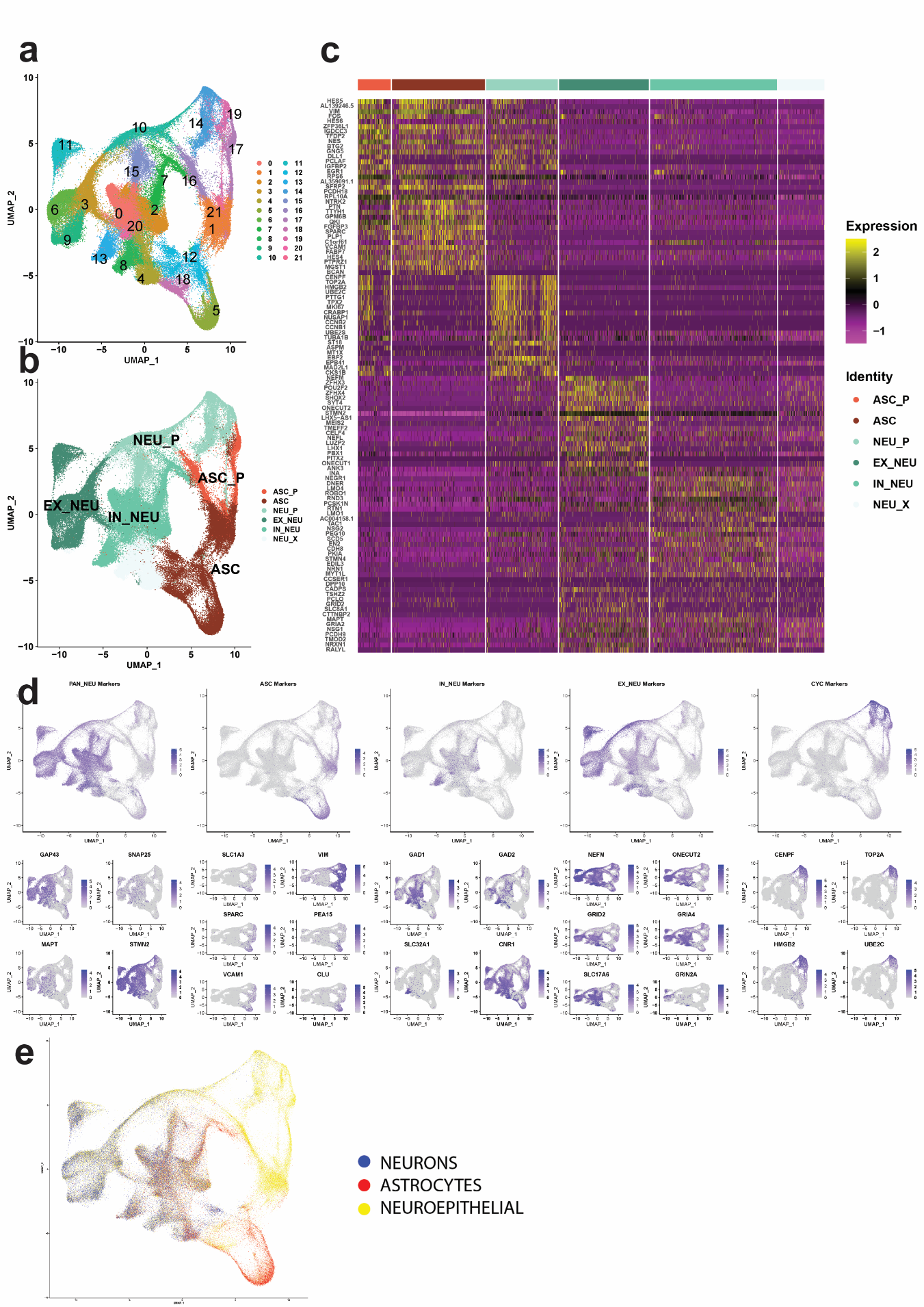

**Supplemental Figure 5. scRNA-seq clustering and cell type identification.**

a. UMAP of unsupervised Seurat clustering based on 30 principle components (PC) producing 21 clusters.
b. UMAP of 130,605 cells from all experimental conditions after filtration and cell type identification (see Methods). Five main cell types are identified, excitatory neurons (EX_NEU, dark green), inhibitory neurons (IN_NEU, light green), neuron precursors (NEU_P, teal), astrocytes (ASC, dark red), and astrocyte precursors (ASC_P, light red).
c. Scaled expression heatmap of the top 20 cell type specific genes for each identified cell type cluster (log fold change > 0.25).
d. UMAP gene expression feature plots of representative canonical cell type markers and an averaged gene expression feature plot of the combined cell type markers for neurons (PAN_NEU), astrocytes (ASC), GABAergic inhibitory neurons (IN_NEU), glutamatergic excitatory neurons (EX_NEU) and cycling neuronal progenitors (CYC) used for manual cell type identification.
e. UMAP of automated cell typing results using the SingleR platform with the Human Atlas fine background (see Methods).

**Supplemental Fig 6**

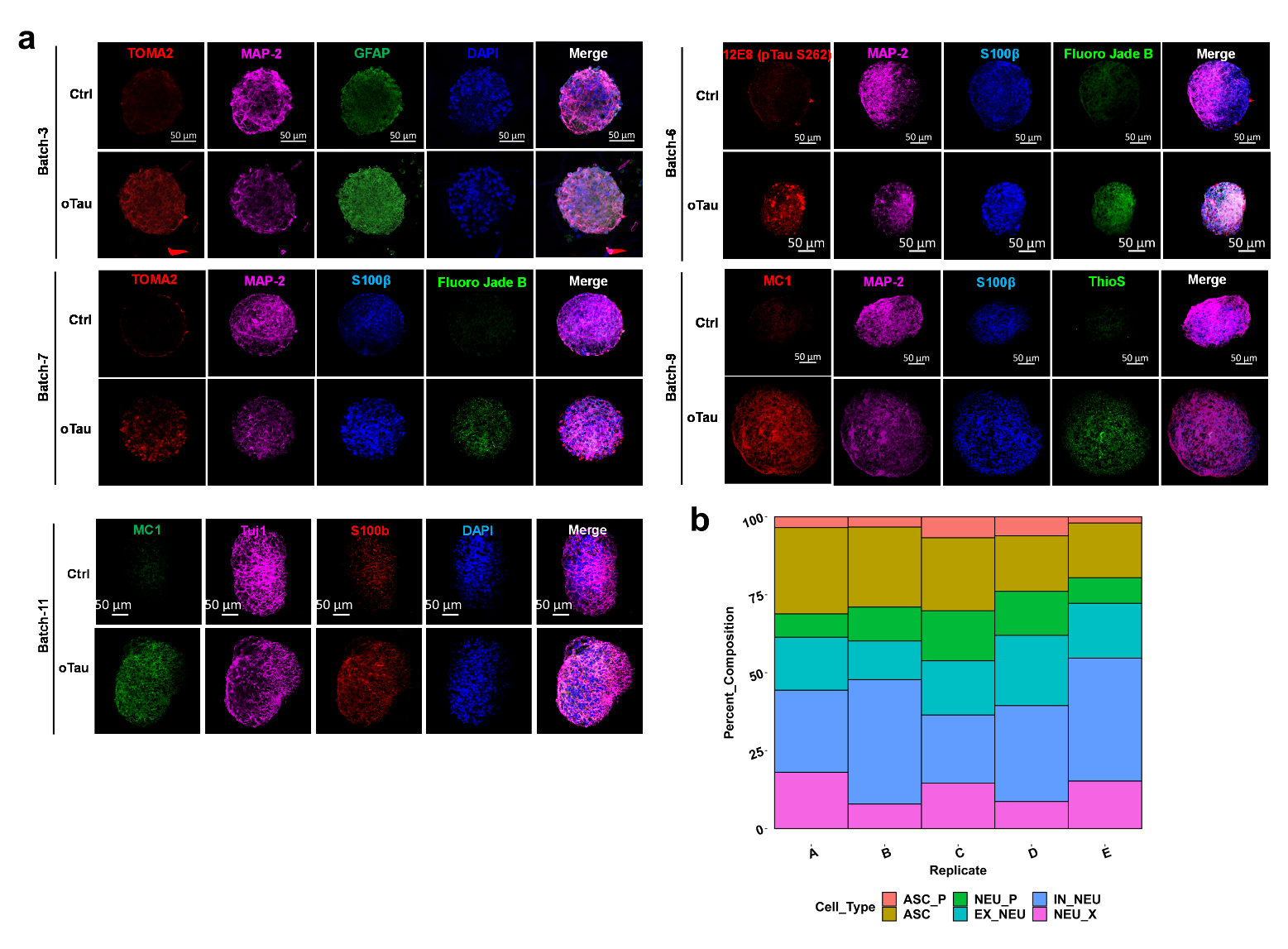

**Supplemental Figure 6. Cell type composition is consistent across replicate batches of asteroids.**

a. Representative images of immuno-fluorescence labeled asteroids from 5 independent batches to show the consistency among different experiments. The experiment condition and the labeling of each fluorescence channel are as indicated in the images. Scale bars = 50 µm.

b. Percent composition of cell types across the 5 replicate asteroid batches by scRNA-seq.

**Supplemental Fig 7**

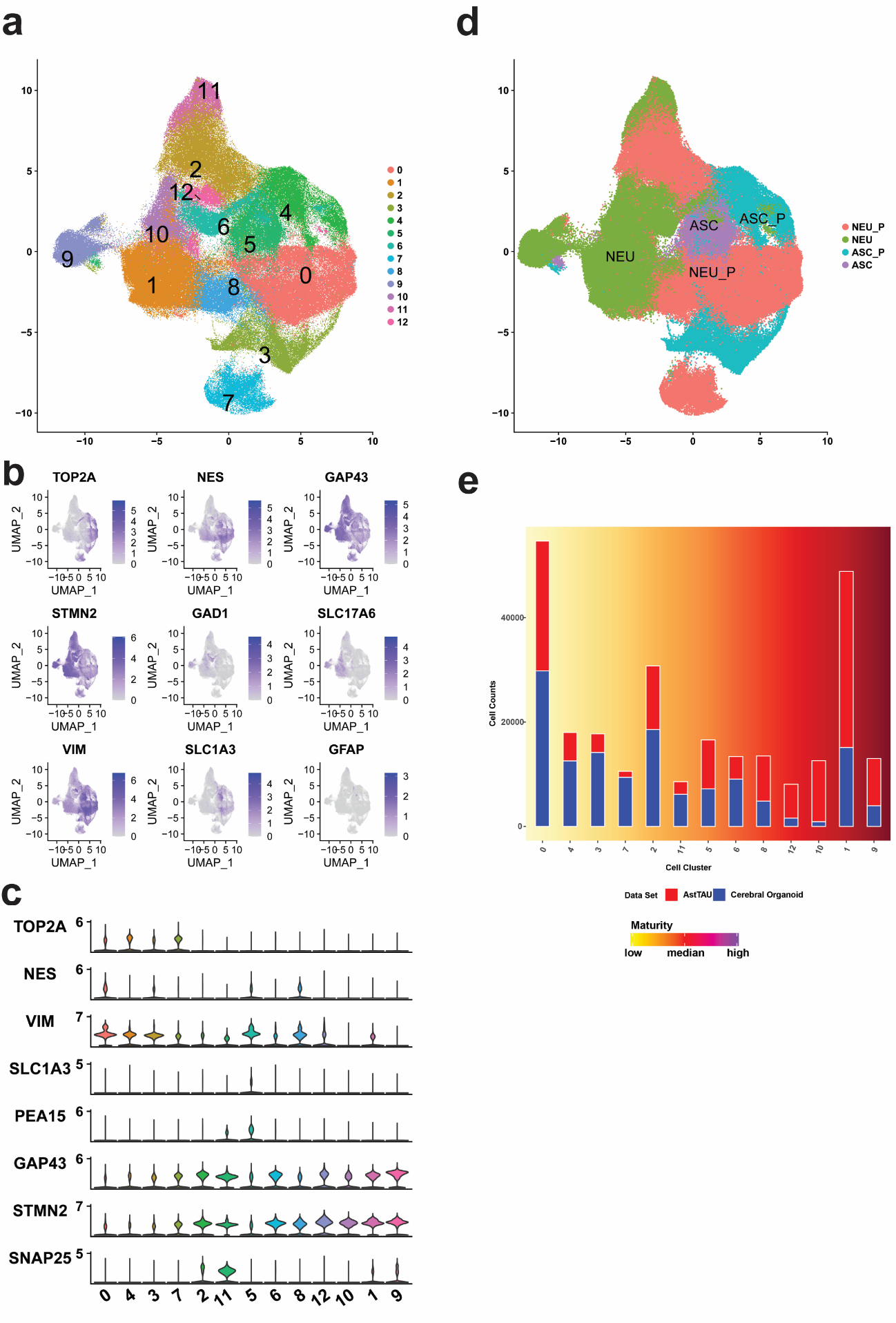

**Supplemental Figure 7.** The AstTau scRNA-seq dataset was reintegrated with scRNA-seq datasets from other 3D cerebral organoid models using the Liger batch correction algorithm (see methods). Clustering, cell typing, and cluster composition analysis was performed on the new integration to compare the maturity of the AstTau model to other 3D cerebral organoid models.

a. UMAP of Liger clustering of the integrated comparative dataset.
b. UMAP gene expression feature plots of representative canonical cell type markers for neurons (GAP43, STMN2), astrocytes (VIM, SLC1A3, GFAP), GABAergic inhibitory neurons (GAD1), glutamatergic excitatory neurons (SLC17A6) and cycling neuronal progenitors (TOP2A, NES) used for manual cell type identification of the integrated comparative dataset.
c. Gene expression of representative canonical cell type markers for neurons (GAP43, STMN2, SNAP25), astrocytes (VIM, SLC1A3, PEA15), and cycling neuronal progenitors (TOP2A, NES) in Liger clusters (Suppl. Fig. 7a), used for manual cell type identification of the integrated comparative dataset.
d. UMAP after cell type identification of the integrated comparative dataset. Four cell types are identified, neurons (NEU, green), neuron precursors (NEU_P, red), astrocytes (ASC, purple), and astrocyte precursors (ASC_P, teal).
e. Over representation of asteroid cell counts (red) in more mature cell clusters (yellow to dark red gradient) in comparison to the comparative cerebral organoid scRNA-seq dataset from Bhaduri et al., 2020 (see Methods).

**Supplemental Fig 8**

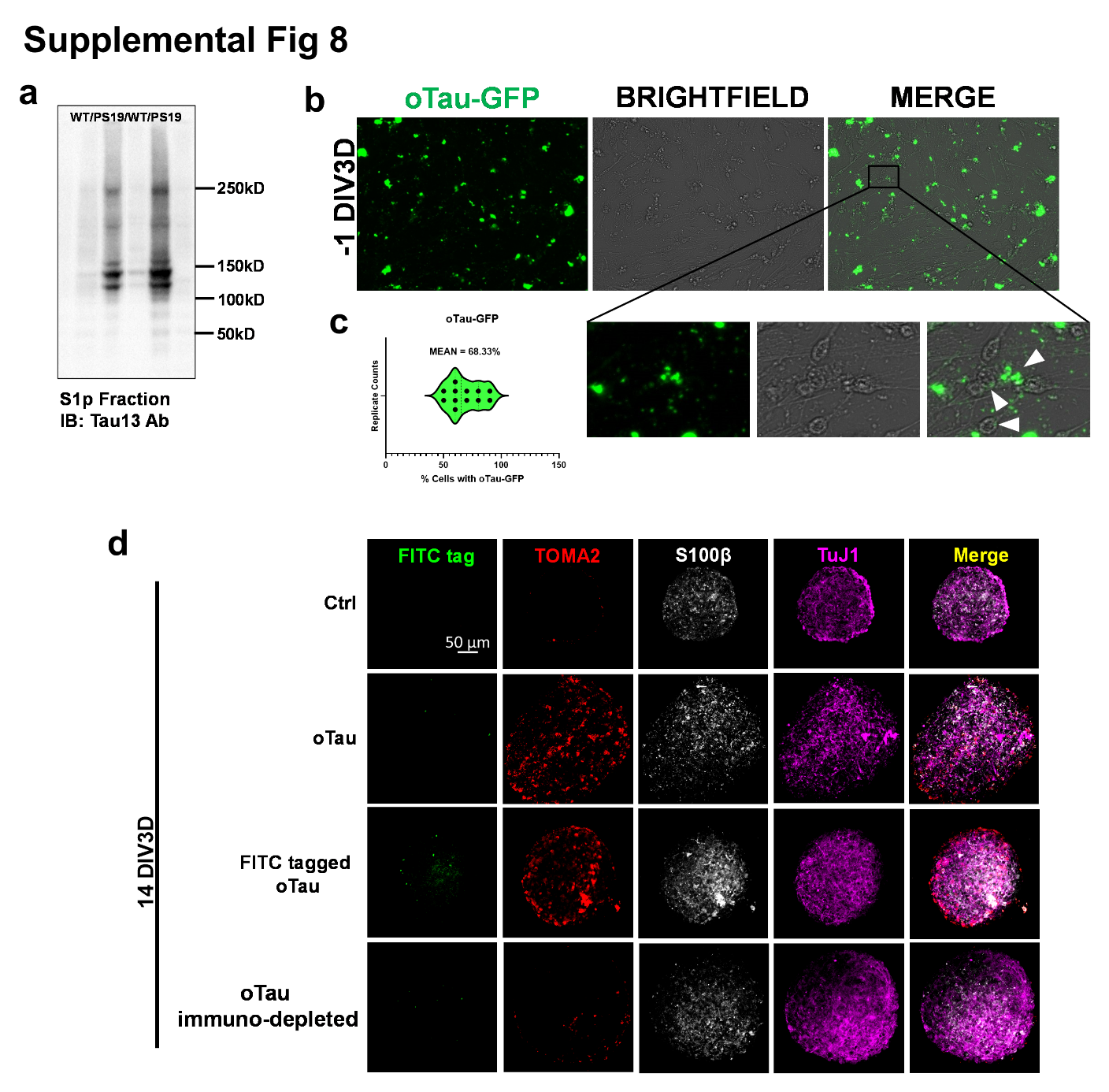

**Supplemental Figure 8. AstTau pathology is specifically induced by oligomeric Tau (oTau).**

a. The molecular weight of oligomeric tau in S1p fractions (oTau) was documented by native page gel electrophoresis. Extracted S1p fraction was quantified by immunoblot with Tau13 antibody and demonstrated the expected oTau bands between 100 kDA and 150 kDA.

b-c. Representative images of brightfield and GFP channels showed the uptake of oTau by hiNCs. S1p oTau fractions were covalently labeled with DyLight 488 (FITC) on amine residues. (Abcam cat# ab201799). hiNCs demonstrated a FITC-tagged oTau uptake efficiency > 68% after 24 hours of exposure, which is the day before 3D asteroid culture establishment.

d. Representative images showing low persistence of FITC-tagged oTau at 14 DIV3D, not correlated with oTau induced toxic oligomer aggregation as marked by TOMA2. Additionally, representative images demonstrating immuno-depleted oTau fraction does not induce toxic oligomer aggregation as marked by TOMA2.

**Supplemental Fig 9**

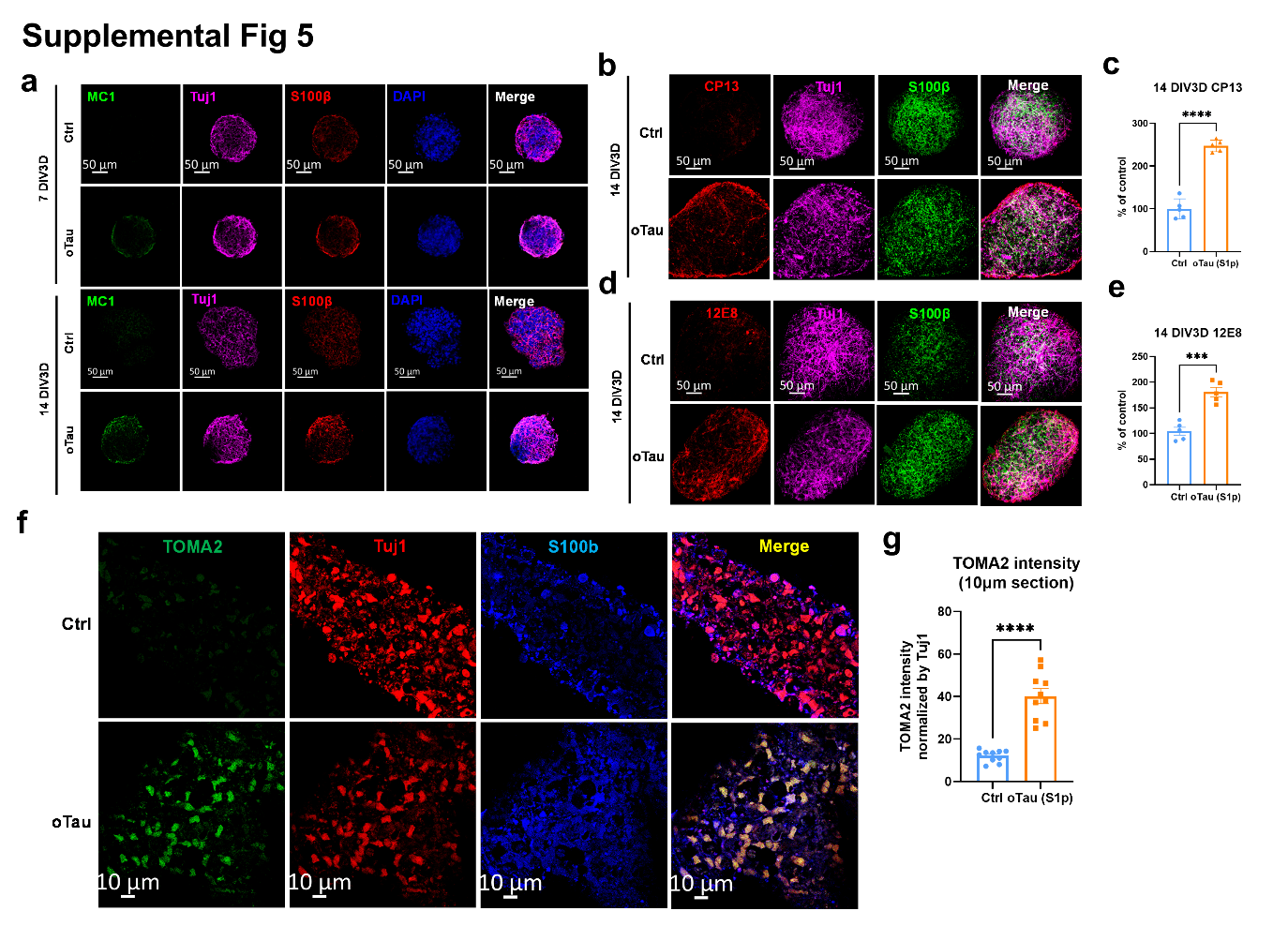

**Supplemental Figure 9. AstTau displays markers of hyperphosphorylated and misfolded tau at 7 and 14 DIV3D.**

a. Representative images showing tau misfolding in oTau seeded asteroids at 7 and 14 DIV3D. The misfolded tau is labeled with MC1 (misfolded tau, green). Neurons were labeled with Tuj1 (βIII tubulin, violet) and astrocyte were labeled with S100β (red). Scale bars = 50 µm.

b. Representative images showing tau hyperphosphorylation in oTau seeded asteroids at 14 DIV3D. Hyperphosphorylated tau is labeled with CP13 (pS202 tau, red). Neurons were labeled by Tuj1 (βIII tubulin, violet) and astrocyte were labeled with S100β (green). Scale bars = 50 µm.

c. Quantification of CP13 labeled fluorescence intensity at 14 DIV3D. Data obtained from 5 independent asteroids. Error bars = SEM. *****p*<0.001 comparisons to vehicle control by two-tailed t test.

d. Representative images showing tau hyperphosphorylation in oTau seeded asteroids at 14 DIV3D. Hyperphosphorylated tau is labeled with 12E8 (pS262 tau, red). Neurons were labeled by Tuj1 (βIII tubulin, violet) and astrocyte were labeled with S100β (green). Scale bars = 50 µm.

e. Quantification of 12E8 labeled fluorescence intensity at 14 DIV3D. Data obtained from 5 independent asteroids. Error bars = SEM. ****p*<0.005 comparisons to vehicle control by two-tailed t test.

f-g. Immuno-fluorescence labelling of 10 µm thin sections of asteroid showed similar labeling patterns as the images shown in figure 2 by integrated asteroids. Scale bars = 20 µm. Quantification of TOMA2 (red) fluorescence intensity was obtained from 10 independent asteroids and normalized by corresponding Tuj1 intensity. Error bars = SEM. *****p*<0.001 comparisons to vehicle control by two-tailed t test.

**Supplemental Fig 10**
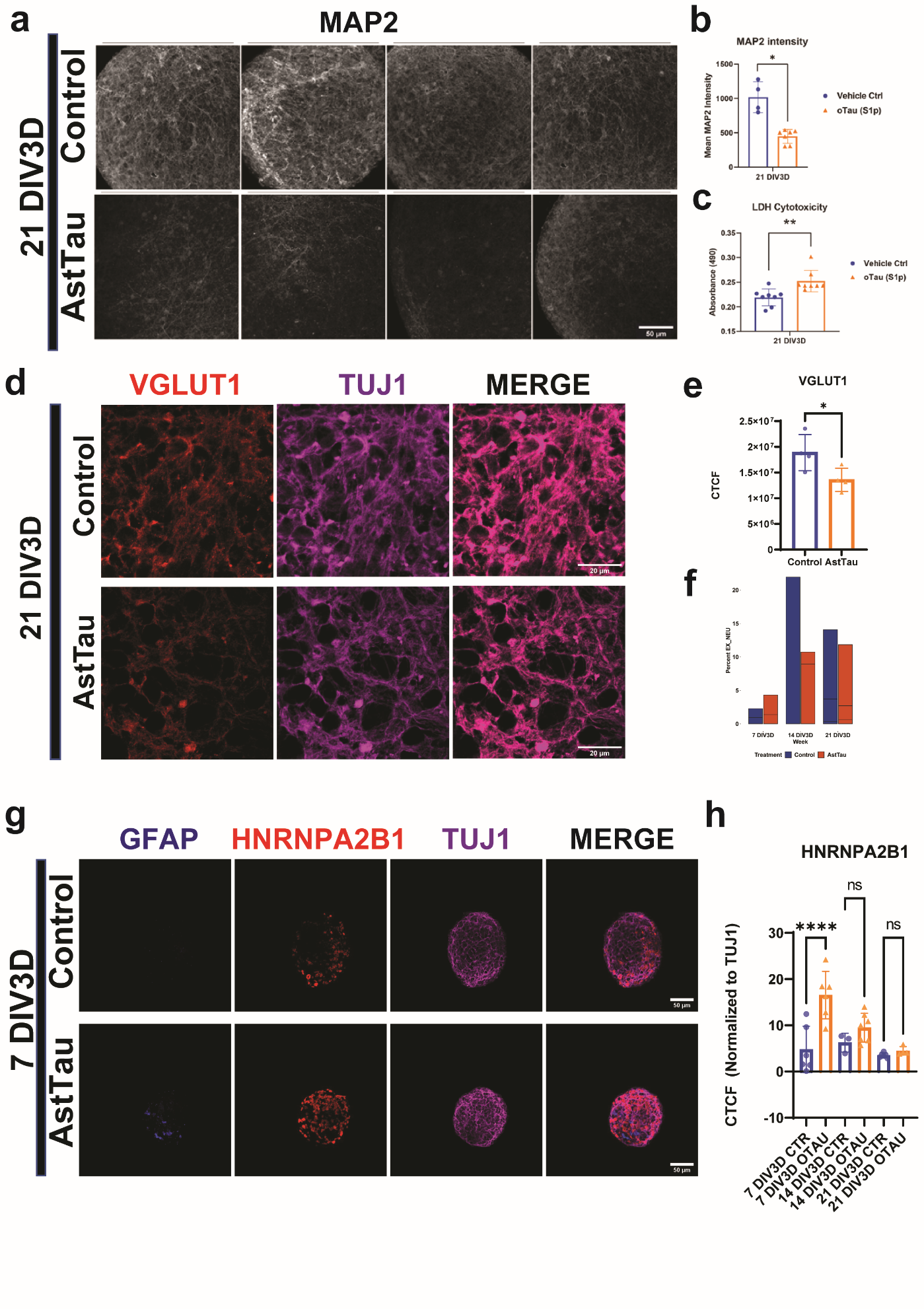

**Supplemental Figure 10. AstTau displays markers of translational stress at 7 DIV3D and neurodegeneration at 21 DIV3D.**

a*.* Quadruplicate representative images of MAP2 immunolabeled control and AstTau at 21 DIV3D demonstrating MAP2 intensity reduction. Scale bars = 50 µm.

b. Quantification of mean MAP2 intensity between control and AstTau. Data obtained from >4 independent asteroid pools. Error bars = SEM. Unpaired t test was performed, *p<0.05.

c. Quantification of increase in cytotoxic LDH release (absorbance 490) between control and AstTau at 21 DIV3D. Data obtained from 8 independent asteroid pools. Error bars = SEM. Unpaired t test was performed, **p < 0.001.

d. Representative images showing glutamatergic synaptic loss in AstTau at 21 DIV3D. Glutamatergic synapses are labeled with VGLUT1 (red) and neurons were labeled with Tuj1 (βIII tubulin, violet) Scale bars = 20 µm.

e. Quantification of corrected total cell florescence of VGLUT1 between control and AstTau. Data obtained from 3 independent asteroids. Error bars = SEM. Unpaired t test was performed, *p<0.05.

f. Percent of excitatory neurons (EX_NEU) across the 7-21 DIV3D time course in control and AstTau indicating excitatory neuron loss at 14 and 21 DIV3D.

g. Representative images showing translational stress granule aggregation in AstTau at 7 DIV3D. Stress granules are labeled with HNRNPA2B1 (red), neurons are labeled with Tuj1 (βIII tubulin, violet), astrocytes are labeled with GFAP (blue). Scale bars = 50 µm.

h. Quantification of corrected total cell florescence of HNRNPA2B1 normalized to Tuj1 between control and AstTau across the 7-21 DIV3D time course. Data obtained from >3 independent asteroids. Error bars = SEM. Two-way ANOVA with Tukey’s multiple comparisons test was performed, ****p<0.001.

**Supplemental Fig 11**

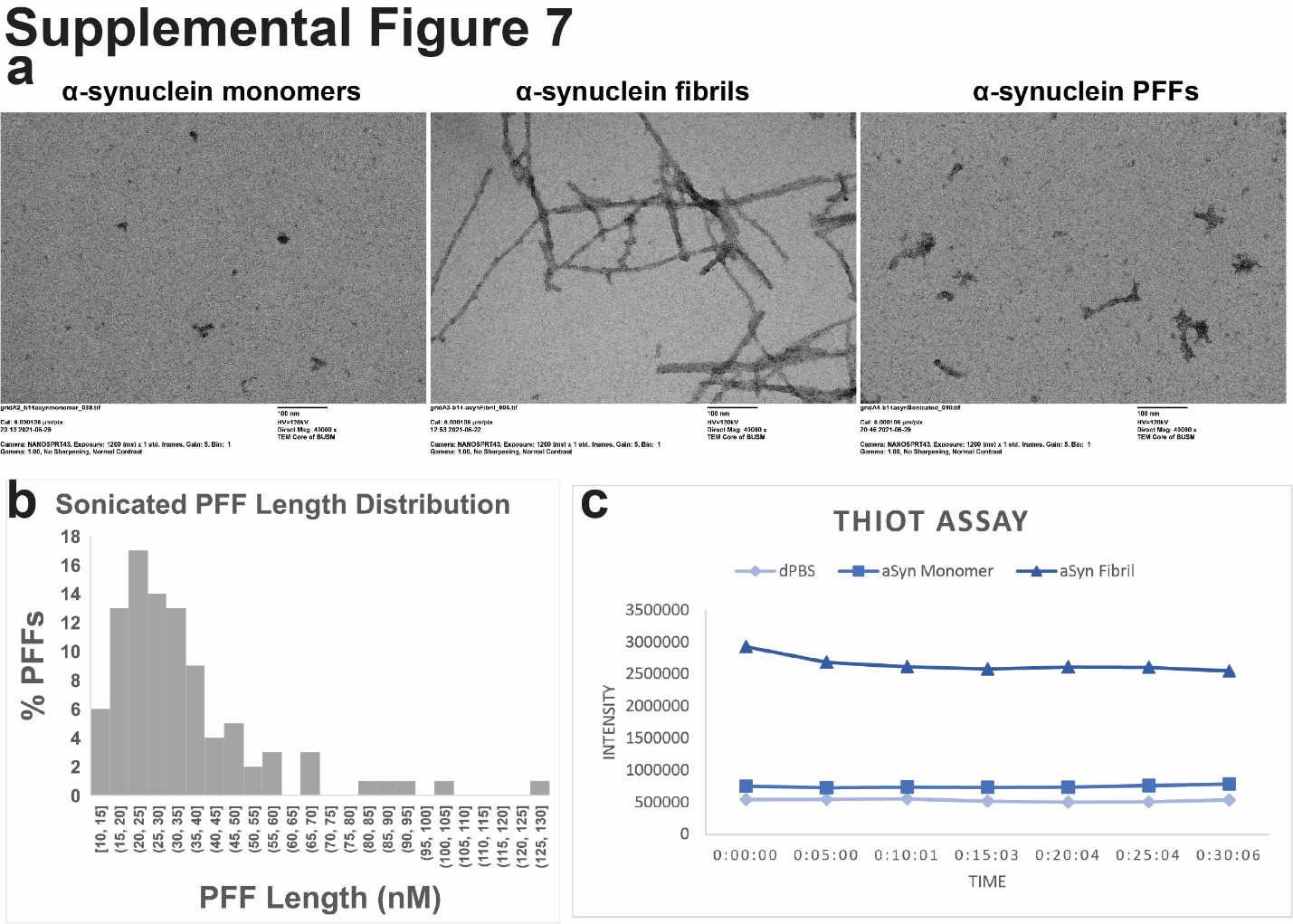

**Supplemental Figure 11. Validation of α-synuclein fibril and PFF formation.**

a. Transmission electron microscopy (TEM) of α-synuclein monomer starting material, prepared α-synuclein fibrils, and sonicated α-synuclein pre-formed fibrils (PFFs). Scale bar= 100 nM (see Method).
b. Quantification of PFF-α-synuclein length distribution used in experimentation.
c. ThioT fibril binding assay validating the presence of prepared α-synuclein fibrils.

**Supplemental Fig 12**

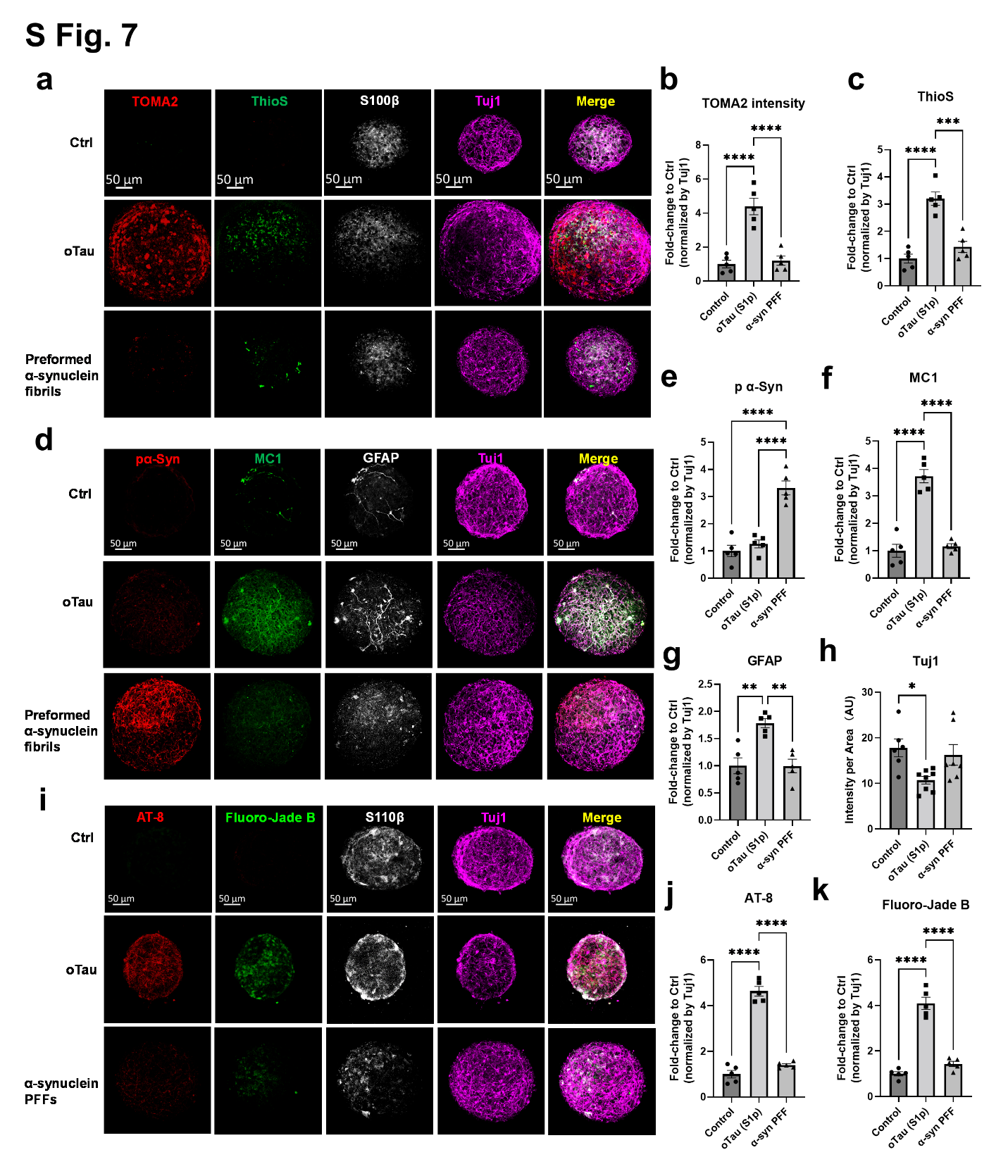

**Supplemental Figure 12. Comparison of pathology and neurodegeneration induced by seeding of oligomeric tau and preformed α-synuclein fibrils, respectively.**

a. Representative images showing tau misfolding in oTau or α-synuclein PFFs seeded asteroids at 21 DIV3D. The oligomeric tau is labeled with TOMA2 antibody (red). The tau fibrils are labeled with Thioflavin S (green). Neurons were labeled with Tuj1 (βIII tubulin, violet) and astrocyte were labeled with S100β (grey). Scale bars = 50 µm.

b-c. Quantification of fluorescence intensity for TOMA2 and ThioS labeled tau aggregates. Data obtained from 5 independent asteroids. Error bars = SEM. *****p*<0.001 and ****p*<0.005 by one-way ANOVA followed with Tukey's multiple comparisons test.

d. Representative images showing tau misfolding and hyperphosphorylated α-synuclein in oTau or α-synuclein PFFs seeded asteroids at 21 DIV3D. The misfolded tau is labeled with MC1 (misfolded tau, green). Phosphorylated α-synuclein is labeled with Anti--synuclein antibody [MJF-R13 (8-8)] (phospho S129, red). Neurons were labeled with Tuj1 (violet) and activated astrocyte were labeled with GFAP (grey). Scale bars = 50 µm.

e. Quantification of phosphorylated α-synuclein fluorescence intensity. Data obtained from 5 independent asteroids. Fluorescence intensities were normalized by Tuj1 intensity and calculated into fold-change of control asteroid. Error bars = SEM. *****p*<0.001 by one-way ANOVA followed with Tukey's multiple comparisons test.

f. Quantification of misfolded tau by MC1 intensity. Data obtained from 5 independent asteroids. Fluorescence intensities were normalized by corresponding Tuj1 intensity and calculated into fold-change of control asteroid. Error bars = SEM. *****p*<0.001 by one-way ANOVA followed with Tukey's multiple comparisons test.

g. Quantification of activated astrocyte by GFAP intensity. Data obtained from 5 independent asteroids. Fluorescence intensities were normalized by corresponding Tuj1 intensity and calculated into fold-change of control asteroid. Error bars = SEM. *****p*<0.001 by one-way ANOVA followed with Tukey's multiple comparisons test.

h. Quantification of neuronal cells by Tuj1 positive intensity. Data obtained from 5-8 independent asteroids. Fluorescence intensities were normalized into intensity per Area (AU). Error bars = SEM. *****p*<0.001 by one-way ANOVA followed with Tukey's multiple comparisons test.

i. Representative images showing tau phosphorylation and neurodegeneration in oTau or α-synuclein PFFs seeded asteroids at 21 DIV3D. The phosphorylated tau is labeled with AT-8 antibody (Ser202/Thr305, red). Degenerating neurons are labeled with Fluoro-Jade B dye (green). Neurons were labeled with Tuj1 (violet), and astrocytes were labeled with S100β (grey). Scale bars = 50 µm.

j-k. Quantification of AT-8 and Fluoro-Jade B intensity respectively. Data obtained from 5 independent asteroids. Fluorescence intensities were normalized by corresponding Tuj1 intensity and calculated into fold-change of control asteroids. Error bars = SEM. *****p*<0.001 by one-way ANOVA followed with Tukey's multiple comparisons test.

**Supplemental Fig 13**

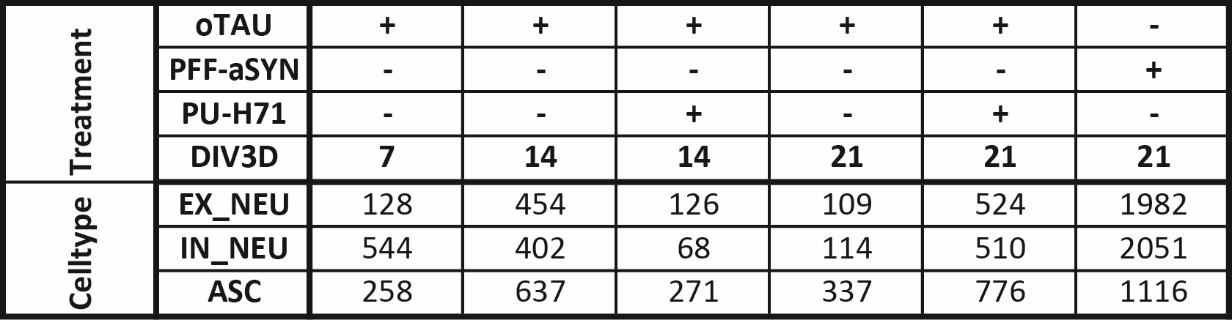

**Supplemental Figure 13. Table of total significant differentially expressed genes counts between control and AstTau by cell type and condition (p< 0.05, log2FC < -0.25 or > 0.25).**

**Supplemental Fig 14**

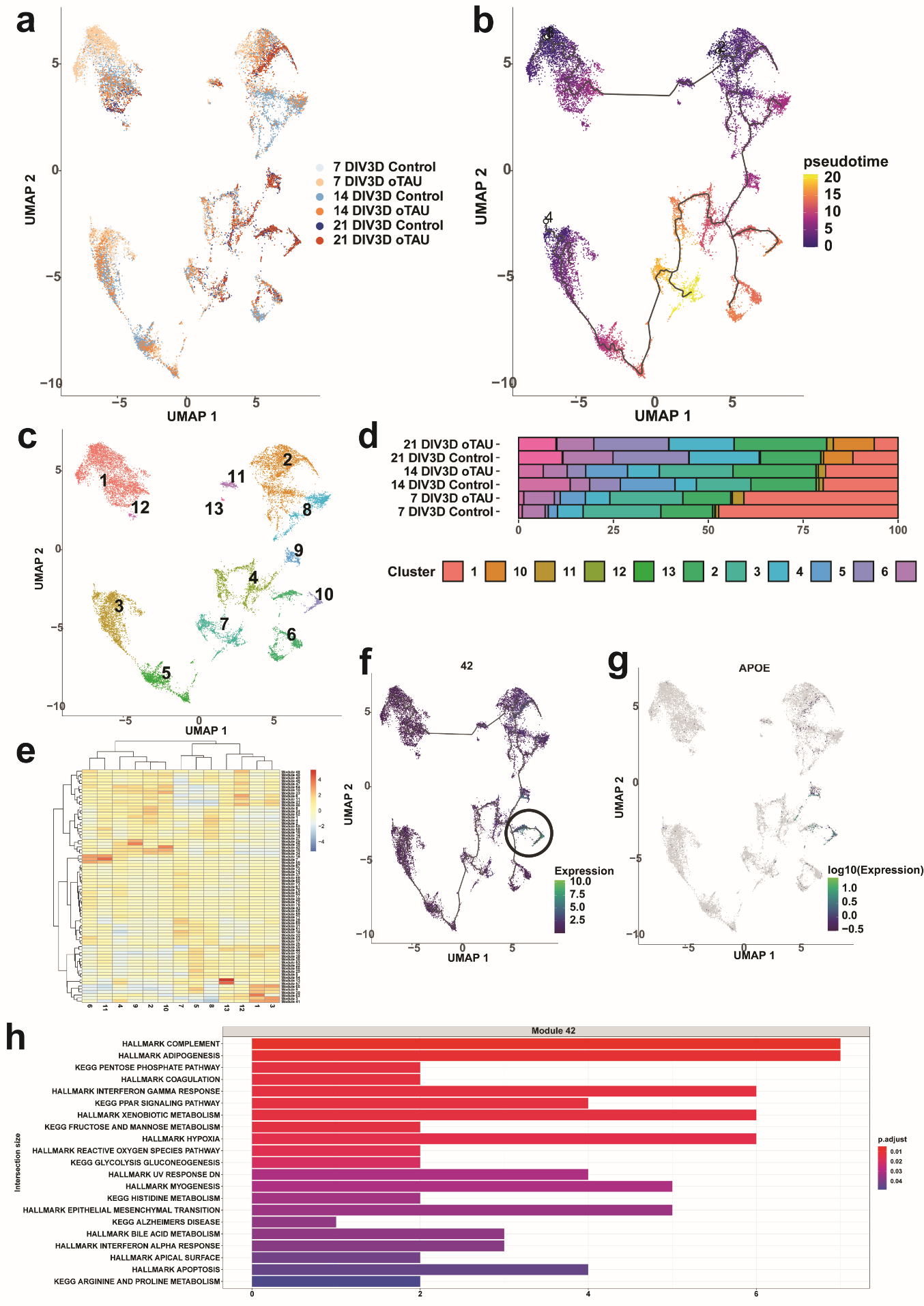

**Supplemental Figure 14. Astrocytes (ASC) were isolated from the scRNA-seq dataset and a pseudotime trajectory was inferred using the Monocle3 package to identify a subset of late stage AstTau enriched ASC and associated gene module signatures.**

a. UMAP of unsupervised Monocle3 clustering of ASC colored by conditional timepoints across the 7-21 DIV3D control and AstTau time course (blue and orange gradients respectively).
b. UMAP of the pseudotime trajectory of ASC, representing the dynamic process of gene expression changes that occur during the progression of AstTau pathology.
c. UMAP of pseudotime clusters 1-12 produced by the Monocle3 clustering.
d. Percent composition of pseudotime clusters in control and AstTau across the 7-21 DIV3D time course revealing an overrepresentation of AstTau ASC
e. in cluster 10.
f. Modules of co-regulated differentially expressed genes within the pseudotime clusters, revealing module 42 upregulated within cluster 10.
g. UMAP expression of module 42 projected on the pseudotime trajectory.
h. APOE expression in the pseudotime trajectory, presenting with module 42 in cluster 10.
i. Top 21 functional gene set enrichment pathways by adjusted p-value of the differentially expressed genes defining module 42 (see Methods).

**Supplemental Fig 15**

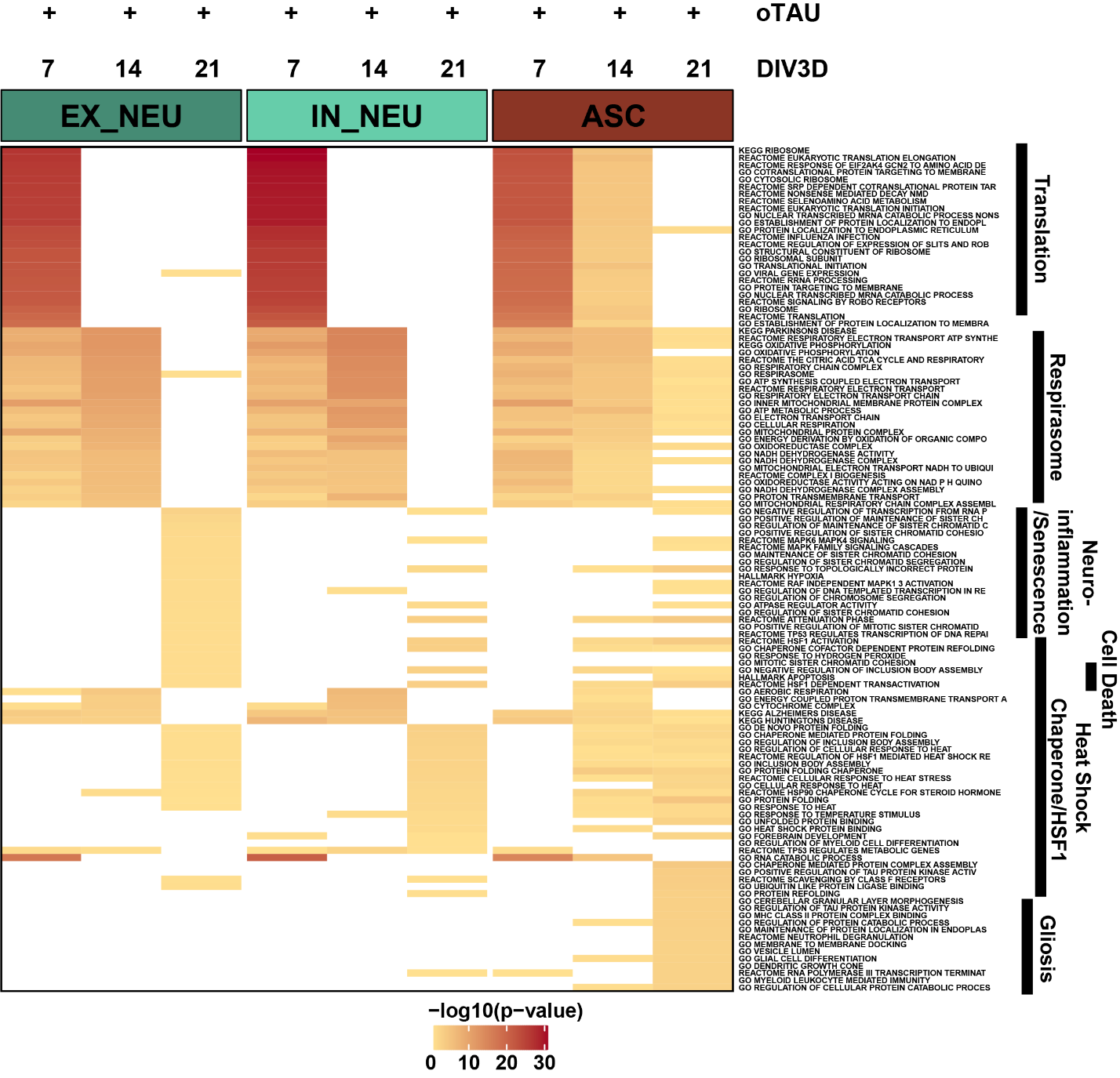
 **Supplemental Figure 15. Top 25 pathways by p-value from functional gene set enrichment of significant differentially expressed upregulated genes (see Methods, p <0.05, fold change > 0.25) between AstTau and control for each timepoint in EX_NEU, IN_NEU, and ASC cell populations presented as –log10 p-value highlighting trends of cell type specific pathway perturbations. Manual annotation of shared gene set features presented for clarity. (See Suppl. SourceTable2.Fig5a7a).**

**Supplemental Fig 16**

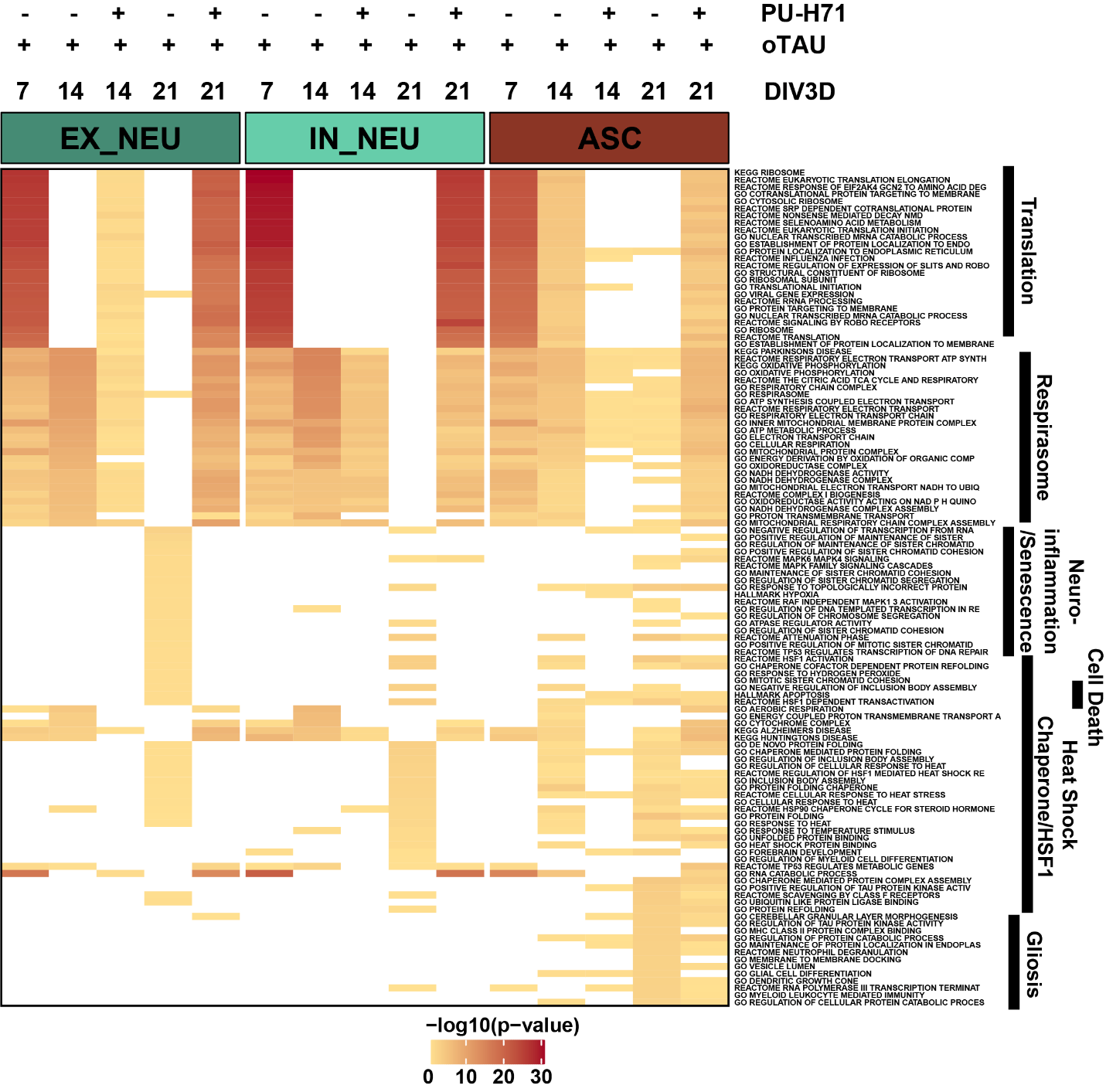

**Supplemental Figure 16. Top 25 pathways by p-value from functional gene set enrichment of significant differentially expressed upregulated genes (see Methods, p <0.05, fold change > 0.25) between AstTau and control for each timepoint in EX_NEU, IN_NEU, and ASC cell populations with and without PU-H71 treatment presented as –log10 p-value highlighting cell type specific pathway perturbations that are ameliorated by PU-H71 treatment. Manual annotation of shared gene set features presented for clarity. (See Suppl. SourceTable2.Fig5a7a).**

**Supplemental Fig 17**

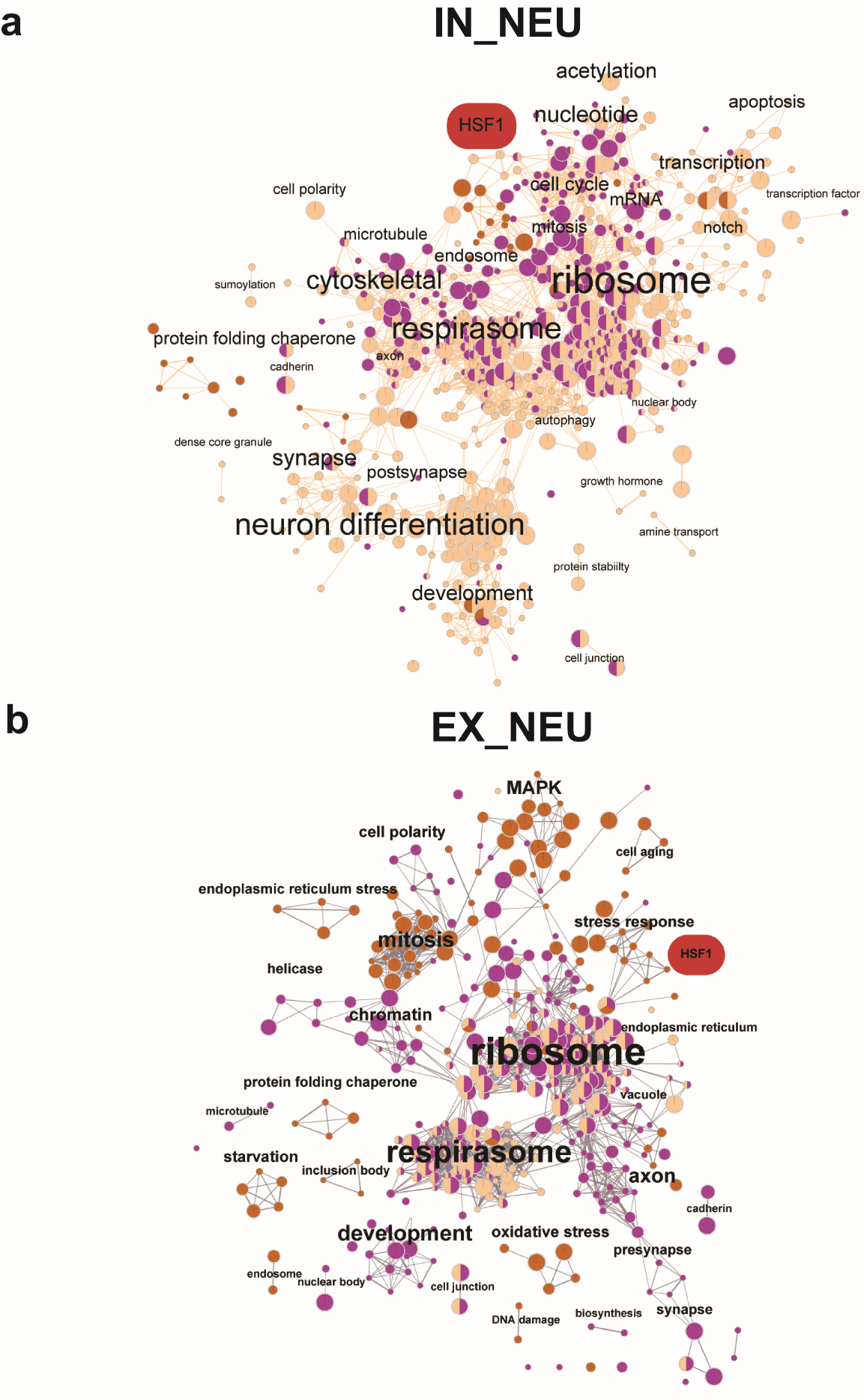

**Supplemental Figure 17. EnrichmentMap clustering presentation of functional gene set enrichment results (see Methods, p < 0.05) with manual annotations across the 7 DIV3D -vehicle (light orange), 21 DIV3D +PU-H71 (purple), and 21 DIV3D-PU-H71 (dark orange) for significantly upregulated DEGs in IN NEU (a) and EX_NEU (b), identifying a range of upregulated responses in AstTau that are ameliorated with PU-H71 treatment. An HSF1 associated cluster is highlighted in red.**

**Supplemental Video 1. Astrocytes display morphological changes in response to oTau induced pathology in AstTau.**

a. 3D projection of a control asteroid at 14 DIV3D at 40X magnification with 6 Z confocal imaging slices captured every 1 M. Nuclei are marked by DAPI (blue), astrocytes are marked by S100β (green), and neurons are marked by TUJ1 (magenta). Scale bars = 50 µm.
b. 3D projection of an AstTau asteroid at 14 DIV3D at 40X magnification with 6 Z confocal imaging slices captured every 1 M. Nuclei are marked by DAPI (blue), astrocytes are marked by S100β (green), and neurons are marked by TUJ1 (magenta). Scale bars = 50 µm.
